# PolarMorphism enables discovery of shared genetic variants across multiple traits from GWAS summary statistics

**DOI:** 10.1101/2022.01.14.476302

**Authors:** Joanna von Berg, Michelle ten Dam, Sander W. van der Laan, Jeroen de Ridder

## Abstract

Pleiotropic SNPs are associated with multiple traits. Such SNPs can help pinpoint biological processes with an effect on multiple traits or point to a shared etiology between traits. We present PolarMorphism, a new method for the identification of pleiotropic SNPs from GWAS summary statistics. PolarMorphism can be readily applied to more than two traits or whole trait domains. PolarMorphism makes use of the fact that trait-specific SNP effect sizes can be seen as Cartesian coordinates and can thus be converted to polar coordinates r (distance from the origin) and theta (angle with the Cartesian x-axis). r describes the overall effect of a SNP, while theta describes the extent to which a SNP is shared. r and theta are used to determine the significance of SNP sharedness, resulting in a p-value per SNP that can be used for further analysis. We apply PolarMorphism to a large collection of publicly available GWAS summary statistics enabling the construction of a pleiotropy network that shows the extent to which traits share SNPs. This network shows how PolarMorphism can be used to gain insight into relationships between traits and trait domains. Furthermore, pathway analysis of the newly discovered pleiotropic SNPs demonstrates that analysis of more than two traits simultaneously yields more biologically relevant results than the combined results of pairwise analysis of the same traits. Finally, we show that PolarMorphism is more efficient and more powerful than previously published methods.

## Introduction

Genetic variation in the genome partly explains phenotypic differences between individuals. Genome-wide association studies (GWAS) aim to identify the specific genetic variants (usually single nucleotide polymorphisms, SNPs) that are associated with phenotypic variation. Over the past decades, GWAS have led to the discovery of thousands of SNP-trait associations [1], [2].

From these discoveries we know that some SNPs can influence multiple traits; i.e. they are pleiotropic [3]. Pleiotropy is widespread in the human genome. An association analysis between millions of SNPs and hundreds of traits found that almost ten percent of SNPs were associated with more than one trait [4]. Moreover, pleiotropic SNPs have been identified for many trait combinations. In many cases, the traits are known to be biologically related; pleiotropic SNPs have been identified for several psychiatric phenotypes [5] and different types of cancers [6]. However, pleiotropic SNPs have also been described for seemingly unrelated diseases; for instance for prostate cancer and type 2 diabetes [7], schizophrenia and Human Immunodeficiency Virus (HIV) infection [8], and Alzheimer’s disease and lung cancer [9]. This could mean that those SNPs are involved in a biological process with a more general function. It could also mean that the studied traits are more biologically related than was previously known and might have a common etiology. Identifying more pleiotropic SNPs can thus transform our current classification of diseases [10].

Pleiotropy analysis can also be useful to identify pleiotropic SNPs in druggable genetic targets, which can help predict adverse treatment effects [10] as well as identify diseases that could be treated with existing drugs [11]. Moreover, pleiotropy can be leveraged for more accurate risk prediction [12]. Finally, methods like Mendelian Randomization (MR) rely on the assumption that there is no direct effect of the SNPs used on both exposure and outcome [13]. Since pleiotropy methods can be used to indicate whether some SNPs are pleiotropic, they can be used to filter these SNPs.

It should be noted that analysis of similarity between traits can also be done using genetic correlation, but this answers a different question. Genetic correlation gives the overall - genome-wide - correlation of effect sizes. Pleiotropic SNPs have a shared effect regardless of the genetic correlation and may tag a specific biological pathway or process rather than describing a general relationship between two traits. If traits are correlated and often co-occur in individuals, then any SNP that affects trait X will also be associated with trait Y, even if it does not directly affect trait Y. These SNPs are not actually pleiotropic because they are only directly associated with one trait. For this reason, to identify pleiotropic SNPs it is not sufficient to take the intersection of SNPs that are associated with both traits. Even if the traits are uncorrelated, intersecting SNP-sets is not an optimal approach; both GWASs need to be sufficiently powered to discover the pleiotropic SNP. Moreover, SNPs that are found with this approach lack an important feature: we know that they are shared but we do not know how shared they are and if this might be statistically significant.

Recently, a few methods that aim to identify pleiotropic SNPs have been described. HOPS [14] and PLEIO [15] both identify a SNP as shared if it is associated with at least one of the traits of interest. Problematically, SNPs with an effect on only one trait will thus also be identified and cannot readily be differentiated from truly pleiotropic SNPs. Two other methods, PLACO [7] and PRIMO [16], identify a SNP as shared if it is associated with all traits of interest. PLACO can only be used for identification of SNPs that are shared by two traits. Moreover, we will show that PLACO has a high computational burden. PRIMO, on the other hand, only identifies a subset of the pleiotropic SNPs that PLACO finds.

Here, we present PolarMorphism, a new approach to identify pleiotropic SNPs that is more efficient, identifies the same number of pleiotropic SNPs as PLACO, but can be applied to more than two traits. This enables the identification of SNPs that have an effect on numerous traits, and possibly play a role in more general biological processes. PolarMorphism is based on a transformation of the trait-specific effect sizes *x* and *y* to polar coordinates *r* (*radius*, the distance from the origin) and ***θ*** (*theta*, the angle with the x-axis). As a result, *r* is a measure of overall effect and ***θ*** a measure of sharedness, which can be used for downstream significance analysis and SNP ranking.

PolarMorphism enables construction of a trait network showing which traits share SNPs. From SNP-specific networks we observe that most SNPs are associated with traits within one trait domain. We find one SNP - rs495828 in the ABO locus - that is associated with traits across 7 trait domains. We show that analysis of more than two traits is more powerful than the intersection of pairwise results of those same traits. We provide PolarMorphism as an R package on Github under the MIT license: https://github.com/UMCUGenetics/PolarMorphism.

## Results

### Defining pleiotropy

Pleiotropy can be identified in different ways ([3], [17] and Figure 1). Horizontal pleiotropic SNPs directly affect multiple traits. Vertical (or mediated) pleiotropic SNPs directly affect one of the traits, but dependence between the traits leads to an association with both traits. The difference between horizontal and vertical pleiotropy is particularly important in the context of Mendelian randomization (MR). With MR, the causal effect of an exposure (e.g. smoking) on an outcome (e.g. lung cancer) can be determined. Genetic variants that are associated with the exposure are used as so-called ‘instrumental variables’. One important assumption is that these variants only have an effect on the outcome through the exposure. In other words, that they are vertically pleiotropic and not horizontally. Horizontally pleiotropic SNPs - which have a direct effect on both smoking and lung cancer - violate this assumption and should therefore not be used as instrumental variables in MR [18]. The final pleiotropy type is spurious pleiotropy, which can arise from bias in measuring association [19]. For example, one marker SNP can be associated with two or more traits due to that marker being in Linkage Disequilibrium (LD) with another SNP that directly affects one of the traits and yet another SNP that directly affects another trait. The marker SNP seems to be pleiotropic, while in reality neither the marker SNP nor the nearby linked SNPs are pleiotropic. Determining whether the same SNP is likely causal for both traits is only possible with colocalization approaches [20]. Another source of spurious pleiotropy is misclassification of traits. If certain symptoms are shared by two diagnoses, individuals with these overlapping symptoms can be given either diagnosis. As a result, the genetic associations for these diagnoses will be highly correlated. Finally, shared controls and ascertainment bias (participant recruitment in a specific disease field) can also cause spurious pleiotropy [21].

**Figure 1.**
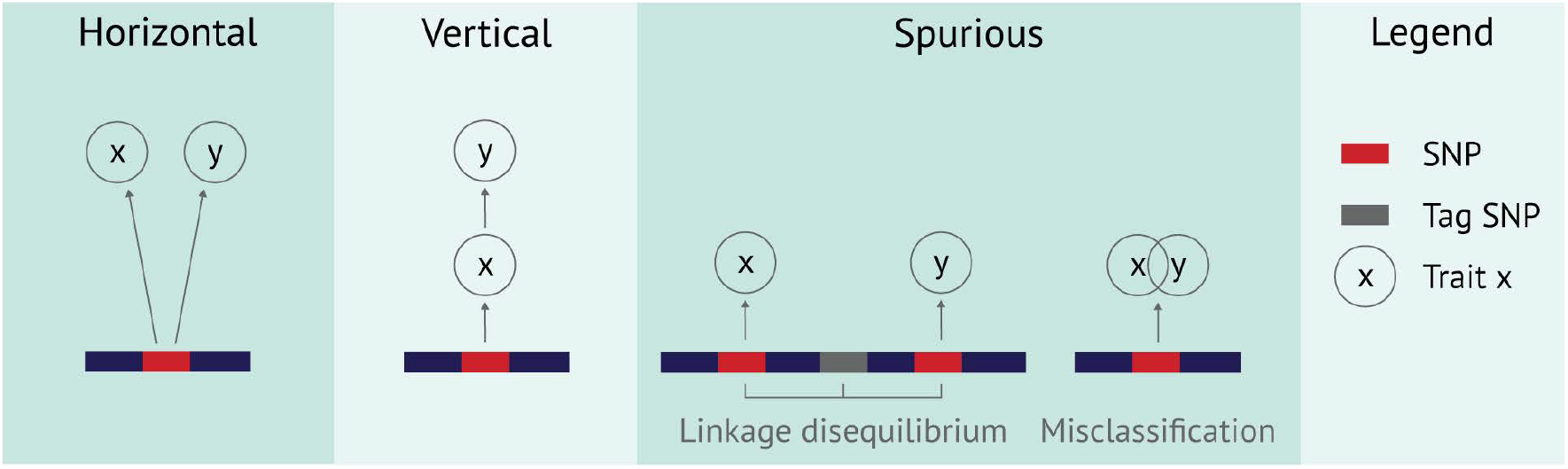
Visualization of horizontal, vertical and spurious pleiotropy, respectively. A horizontally pleiotropic SNP has an effect on all traits under consideration. A vertically pleiotropic SNP has an effect on only one of the traits, but because the traits are correlated it is also associated with the other trait. A SNP can seem pleiotropic because it is in linkage disequilibrium with two SNPs that each individually have an effect on a trait. Misclassification of individuals can also give rise to a seemingly pleiotropic effect.

### Overview of PolarMorphism

We aim to identify pleiotropic SNPs from GWAS summary statistics using an approach that can be routinely applied to combinations of two or more traits. After obtaining summary statistics with effect size beta and standard error SE, we calculate z-scores (beta / SE) per SNP. PolarMorphism can be applied on any number of traits, but here we explain the application to two traits. Analyzing more than two traits requires a slightly different approach (see the methods for a full description) but leverages the same principles.

Our aim is to identify horizontally pleiotropic SNPs. Therefore we first perform a decorrelating transform to attenuate vertical pleiotropy resulting from genetic correlation. Given summary statistics for trait x and y, we calculate a covariance matrix, and use this to apply decorrelation or whitening (see methods for details) yielding decorrelated summary statistic vectors 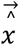 and 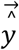. Next the trait-specific vectors 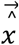 and 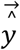 are used to calculate polar coordinates *r* (the distance from the origin) and ***θ*** (the angle with the x-axis, ranging from 0 to 2π). For SNPs that are specific to trait X, ***θ*** is close to 0 or π. For SNPs that are specific to trait Y, ***θ*** is close to 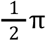 or 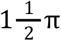. For SNPs that are shared, ***θ*** is approximately 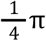 or 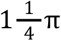 for concordant direction of effect and 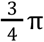 or 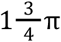 for opposite direction of effect. Each quadrant of the x,y plot only differs in direction of effect in the original GWAS. To simplify further analysis we use the fourfold transform of *θ* (***θ_trans_***), which folds the quadrants on top of each other (equivalent to using the absolute values of the z-scores) and then stretches the angles so they still describe a full circle (Figure 2).

**Figure 2.**
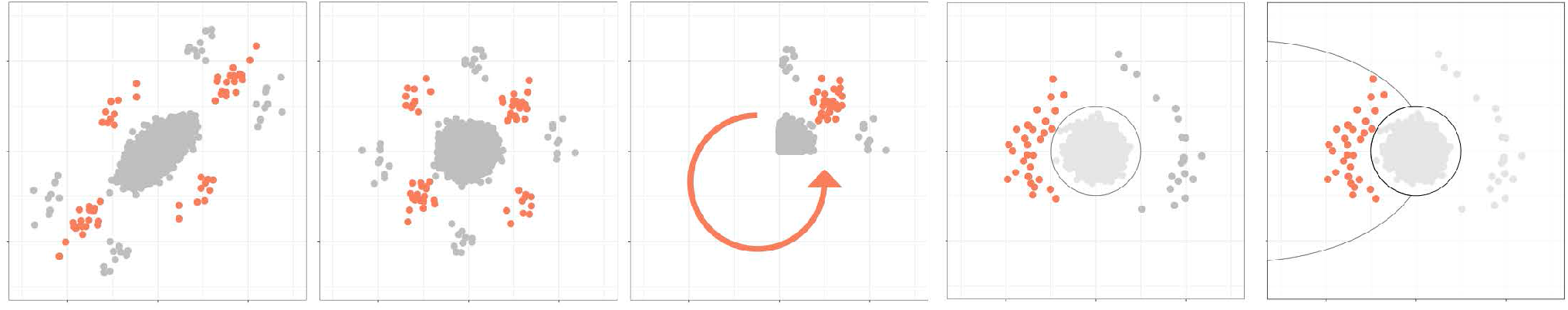
Overview of the method for 2 traits. Z-scores for each trait are plotted on each axis and the data is decorrelated. Cartesian coordinates are transformed to polar coordinates. The absolute values of the z-scores are calculated and the angle is multiplied by 4. After subsetting on SNPs with a significant distance, we calculate p-values for the angle.

To assess significance of sharedness, we separately test the distance *r* and angle ***θ***. Under a null hypothesis of no overall effect, *r* is the square root of a sum of squared normally distributed variables with mean 0. We thus use a central chi distribution to calculate p-values for *r*. Under a null hypothesis of trait-specific effect, ***θ_trans_*** is equal to 0. To calculate p-values for ***θ_trans_*** we use a von Mises distribution with concentration parameter κ. We show that κ depends on *r* (see methods). Estimates of κ from simulations under the null hypothesis are included in the R package. These are used to establish one p-value per SNP.

### Inferring relationships between traits from pleiotropic SNPs

We applied PolarMorphism to all pairwise combinations of 41 traits from different trait domains (table 1). The resulting pleiotropy network is shown in Figure 3. Herein, traits are nodes and the edge weights indicate the number of pleiotropic SNPs discovered by PolarMorphism. The contribution of each SNP to the edge weights is weighted by the inverse of the total number of traits it is associated with, in order to account for the effect that SNPs affecting many traits probably tag a biological process with a general function. Sharing such a SNP is less meaningful than sharing a SNP with an effect on only some traits.

**Figure 3.**
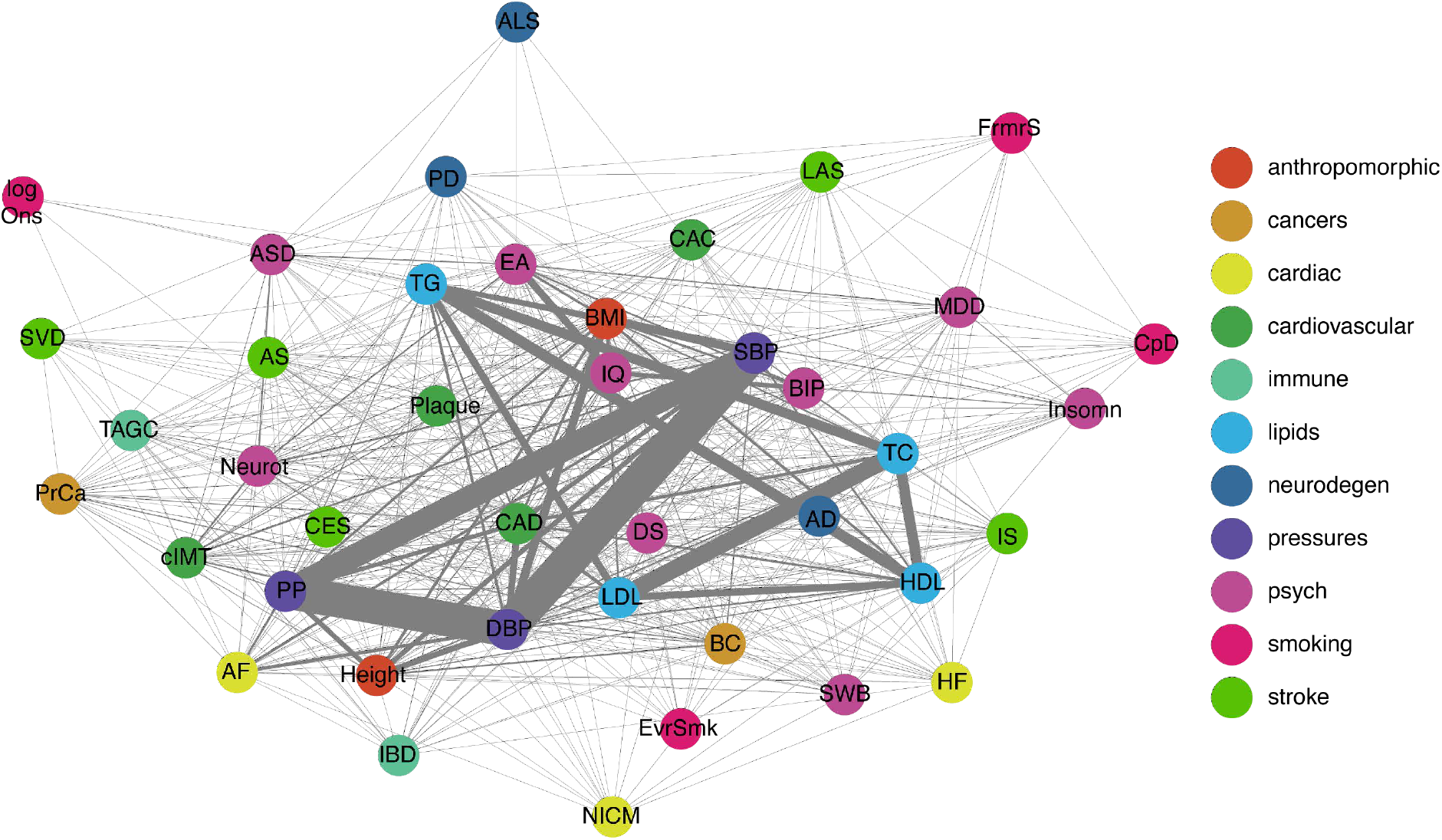
Trait network based on pleiotropic SNPs. The thickness of the lines (network edges) indicates how many loci are shared between two traits (network nodes). Colored by disease domain.

**Table 1.** Trait domains and trait abbreviation as used in the figures.

| Domain name | trait abbreviation | trait name |
| --- | --- | --- |
| anthropomorphic | BMI | body mass index |
|  | Height | height |
| cancers | PrCa | prostate cancer |
|  | BC | breast cancer |
| cardiac traits | AF | atrial fibrillation |
|  | HF | heart failure |
|  | NICM | non-ischemic cardiomyopathy |
| cardiovascular | CAC | coronary artery calcification |
|  | CAD | coronary artery disease |
|  | cIMT | carotid intima-media thickness |
|  | Plaque | presence of carotid plaque |
| immune | IBD | irritable bowel disease |
|  | Asthma | asthma |
| lipids | HDL | high-density lipoprotein |
|  | LDL | low-density lipoprotein |
|  | TC | triglycerides |
|  | TG | total cholesterol |
| neurodegenerative disease | AD | Alzheimer's disease |
|  | ALS | Amyotrophic lateral sclerosis |
|  | PD | Parkinson's disease |
| pressures | DBP | Diastolic blood pressure |
|  | SBP | Systolic blood pressure |
|  | PP | Pulse pressure |
| psychiatric / psychological | ASD | Autism spectrum disorder |
|  | BIP | bipolar disorder |
|  | DS | depressive symptoms |
|  | EA | educational attainment |
|  | IQ | intelligence quotient |
|  | MDD | major depressive disorder |
|  | Neuroticism | neuroticism |
|  | SWB | subjective well being |
|  | Insomnia | insomnia |
| smoking | EvrSmk | ever smoker |
|  | FmrSmk | former smoker |
|  | logOnset | log of age at onset of smoking |
|  | CpD | cigarettes per day |
| stroke | AS | any stroke (hemorrhagic or ischemic) |
|  | IS | ischemic stroke |
|  | CES | cardio-embolic stroke |
|  | LAS | large artery stroke |
|  | SVS | small vessel stroke |

The resulting pleiotropy network is densely connected (512 out of 820 possible edges), supporting earlier descriptions of widely occurring pleiotropy among traits [4], [21] The lipid domain (HDL, LDL, TG and TC) and blood pressure domain (DPB, SBP and PP) each form a fully connected subgraph. SBP has the highest number of edges (degree), sharing SNPs with 37 of the 41 traits. ALS, which shares SNPs with 5 traits, has the lowest degree. Global analysis of the pleiotropy network thus readily reveals general characteristics of traits and trait domains.

Analyzing the pleiotropy network in more detail, we find that most SNPs are associated with traits within one or across two trait domains (51% and 43%, respectively). We observe one SNP that is associated with traits across 7 trait domains: rs495828, a SNP in the ABO gene, which is ubiquitously expressed across many tissues and cell types [22]. For each trait domain, we determine how many SNPs only have associations within that domain (we call these single domain SNPs), and calculate the percentage of the total number of SNPs that were identified for that domain. We find that the psychiatric traits have the highest percentage of single domain SNPs; one third of all SNPs that are shared with a psychiatric trait are only associated with psychiatric traits. The smoking traits have the lowest percentage of single domain SNPs, suggesting that most smoking-associated variants tag general biological processes rather than smoking-specific processes.

#### A comparison with genetic correlation

Genetic correlation 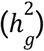 is the correlation of SNP effect sizes on two traits [19]. Non-biological factors like sample overlap between the two GWAS can inflate the 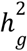 estimate. LDSC [23] or HDL [24] can be used to obtain an 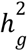 estimate that is not biased by sample overlap. Genetic correlation leads to overall correlation of effect sizes, also in those SNPs with no effect on any of the traits. SNPs that do have an effect can influence 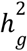 estimates; if they are very pleiotropic they can inflate 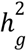, and if they are very trait-specific they can deflate 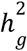. Therefore it is generally recommended to only use the subset of SNPs with no effect on any of the traits for 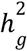 estimation. Also note that pleiotropic effects between traits can be present without genetic correlation, as pleiotropy is a SNP-specific metric and genetic correlation is a genome-wide metric [23].

To assess whether genetic correlation provides the same insight into trait relationships as pleiotropy, we built a network based on genetic correlation. The resulting network is sparse (138 out of 780 possible edges) and only partially overlaps with the pleiotropy network. Figure 4 shows separate subnetworks for edges that exist in both the genetic correlation network and the pleiotropy network or in only one of the two. In total, 416 trait pairs share at least one pleiotropic SNP, but are not genetically correlated (figure 4a). This situation can arise if there are only a few SNPs that are shared but the rest of the genetic architecture of the traits is independent. It is also possible that some shared SNPs have the same direction of effect in both traits while other shared SNPs have an opposite direction of effect, thereby averaging out 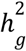. 7 trait pairs are genetically correlated, but do not share any SNPs that are horizontally pleiotropic (figure 4b). Each SNP that is associated with one of the traits is more likely to also be associated with the other, because of the overall 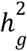 [18]. After decorrelation, vertically pleiotropic SNPs will not be identified by PolarMorphism. 96 trait pairs are genetically correlated and share horizontally pleiotropic SNPs (figure 4c). These are traits that share a number of vertically pleiotropic SNPs, leading to a higher 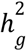, as well as some horizontally pleiotropic SNPs. Our results seem to indicate that two traits are more likely to share at least one pleiotropic SNP than they are to be genetically correlated.

**Figure 4.**
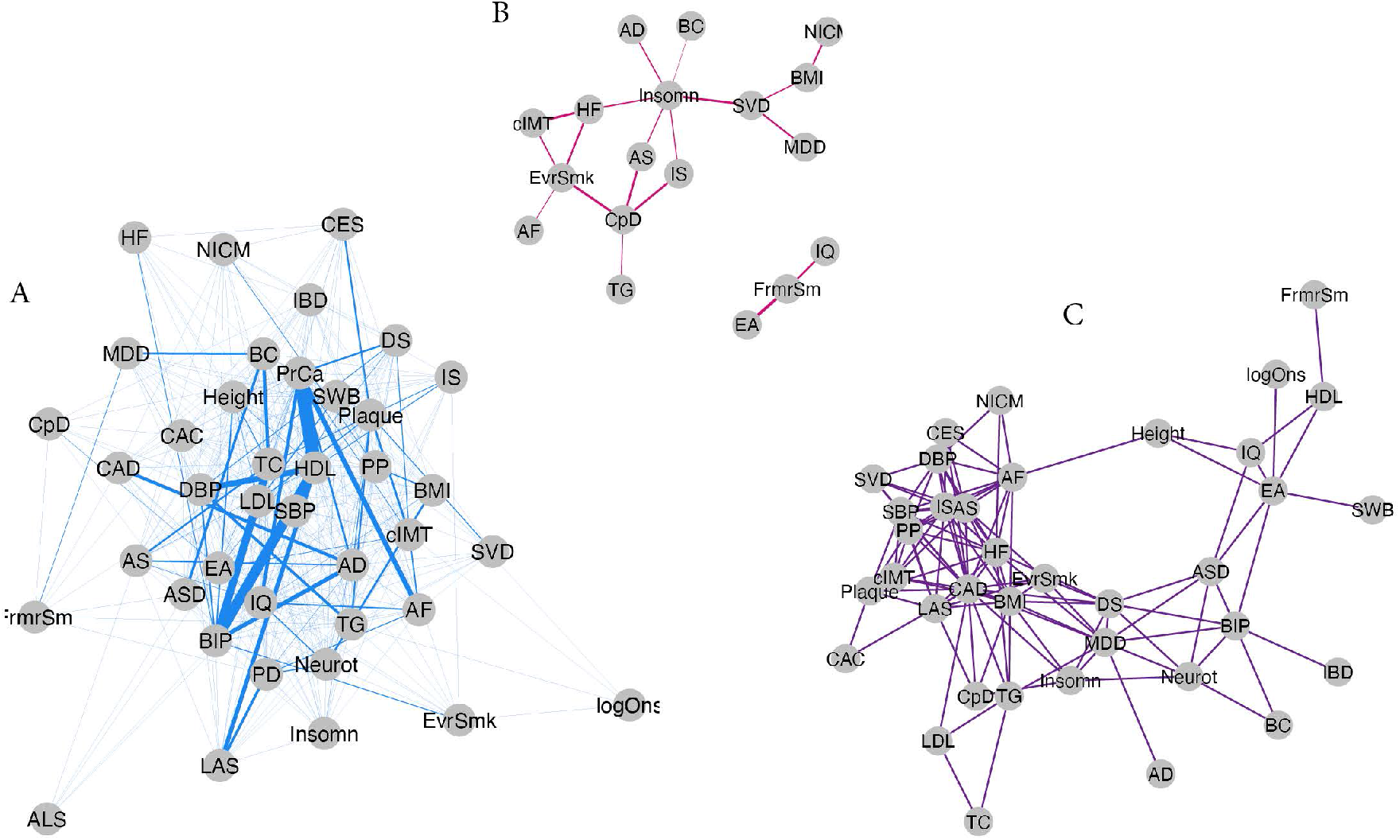
a) Edges denote trait pairs that share pleiotropic SNPs but are not genetically correlated. b) Edges denote trait pairs that are genetically correlated but do not share pleiotropic SNPs. c) Edges denote trait pairs that are genetically correlated and share pleiotropic SNPs.

### The stroke domain

The stroke domain consists of any stroke (AS); its subtype ischemic stroke (IS); and its subtypes cardioembolic stroke (CES), large artery stroke (LAS) and small vessel stroke (SVS). The three IS subtypes are generally believed to have different etiologies [25]–[27], and previous efforts have resulted in tens of subtype-specific associations [28]–[31]. In line with this, our analysis does not reveal any shared SNPs. It should be noted that shared SNPs have been described before for LAS and SVS and for LAS and CES [28]. However, SNPs at these loci were low-confidence and therefore not included in our analysis (see methods for details).

Given the lack of shared SNPs among the IS subtypes, we investigated which other traits share SNPs with each of the IS subtypes. To that end we looked at the subnetwork composed of the IS subtypes and their direct neighbors (figure 5). Our analysis reveals that six traits (CAD, DBP, Plaque, PP SBP, TC) share SNPs with all IS subtypes. This indicates that all ischemic stroke subtypes are associated with biological pathways with a possible effect on blood pressure and lipids. CES shares most pleiotropic SNPs with atrial fibrillation (AF), which is believed to be its main cause [26]. LAS, which is thought to arise from atherosclerotic plaques in the carotid arteries that rupture or block blood flow [31], shares most SNPs with cIMT - a proxy for the extent of carotid atherosclerosis. SVS, which is thought to have a cardiovascular origin like the other IS subtypes [32], shares most SNPs with CAD. Notably, it also shares many SNPs with Alzheimer’s and Parkinson’s disease. This might indicate that many of the SNPs that are associated with risk of small vessel stroke also influence risk of neurodegenerative disease. Note that the edges LAS-HDL, SVS-AD, SVS-PD and SVS-Plaque were only found in the pleiotropy network and not in the genetic correlation network. This indicates that pleiotropic SNPs harbor information that is complementary to genome-wide correlation measures. Furthermore, zooming in on one trait domain shows how PolarMorphism can be employed to gain more detailed insight in trait relationships than the general patterns that can be gathered from the complete network.

**Figure 5.**
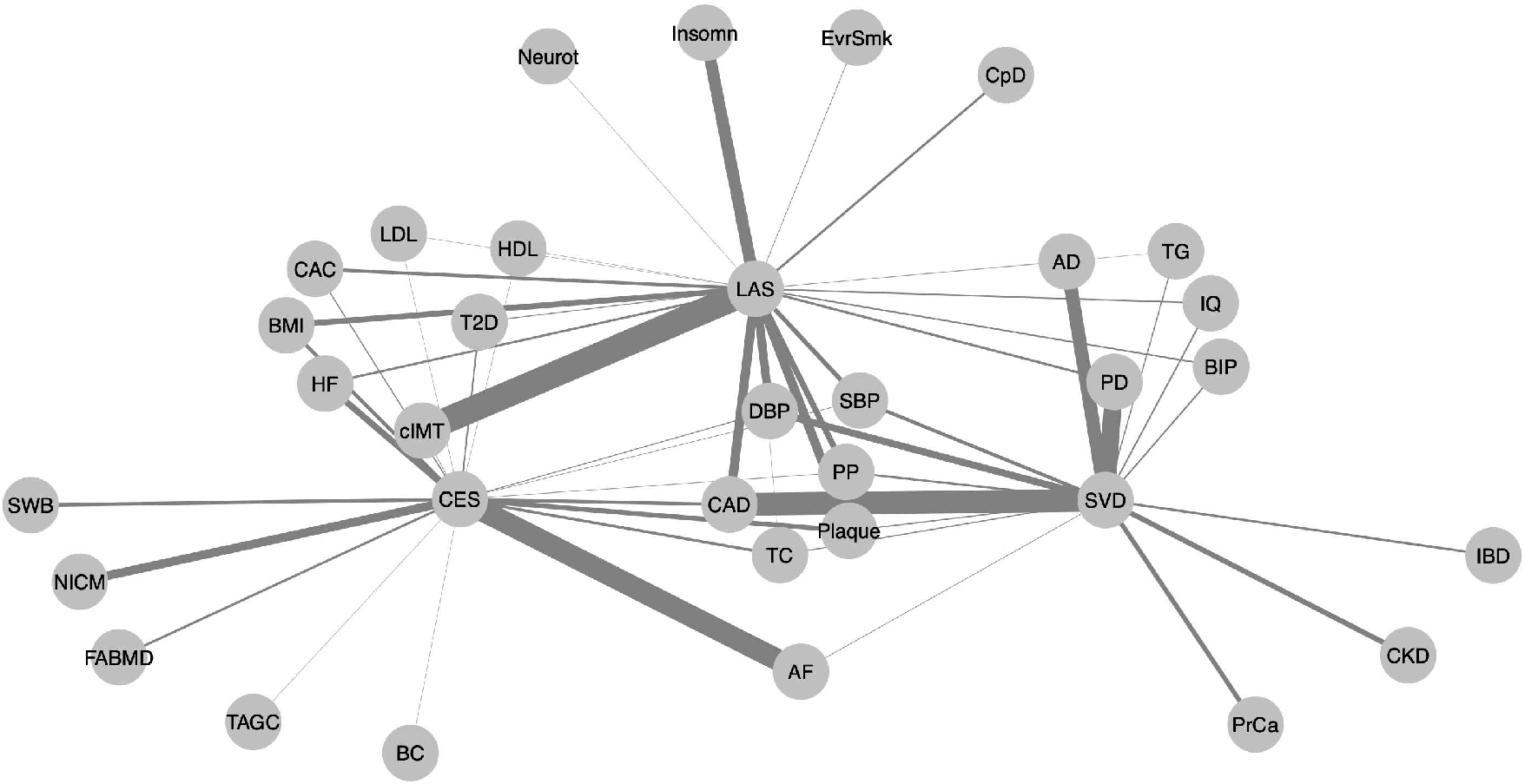
Trait network of the IS subtypes and their direct neighbors, based on the weighted full network as described earlier. Only edges between any of the IS subtypes and any other trait are drawn; in other words, edges between two nodes shown here that do not include an IS subtype, are not drawn.

**Figure 5.**
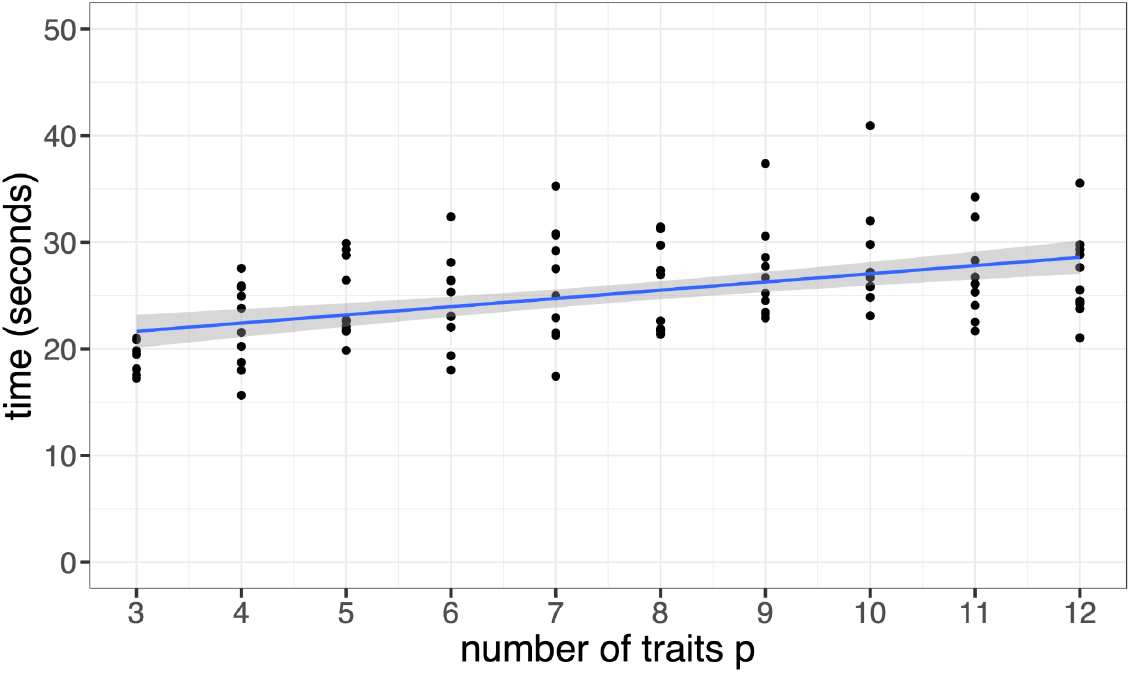
Runtime scales with the number of traits p. a) p ranges from 3 to 12. The slope of the regression line is 0.75 (se = 0.13). b) p ranges from 2 to 12. The large difference between 2 and more than two traits is due to the different methodology that we use if p > 2, see the methods for details.

### Joint analysis of more than two traits identifies more pleiotropic SNPs than pairwise analyses of the same traits

PolarMorphism can be used to find SNPs that are shared by any number of traits. A SNP with a small effect on each trait might not be identified in univariate or even pairwise analysis, but could be if more traits are included. We therefore investigated whether analysis of three or more traits is indeed more powerful than the combined results from pairwise analyses of those same traits. Pairwise analyses of the lipid domain (HDL, LDL, TC, TG) identifies 186 shared loci. Analysis of all four traits together identifies 1029 shared loci. 180 loci are found by both approaches.

To explore whether the increased number of loci is biologically relevant, we perform gene set enrichment analysis in DEPICT [33] on the significant loci from the pairwise analyses and the significant loci from the joint analysis. In order to get the relevant genes for each locus, we perform clumping using DEPICT’s default settings. Hence the number of DEPICT loci differs from the loci that we identified (108 pairwise loci, 496 joint loci, see tables S4 and S6). The pairwise results are enriched for 12 gene sets (table S5) whereas the joint results are enriched for 85 gene sets (table S7). Moreover, the loci revealed by the joint analysis result in enrichments that are more significant: 85 of the 95 gene sets that are significant in either analysis are more significant in the joint analysis, and 2 of the 2 gene sets that are significant in both analyses are more significant in the joint analysis. Moreover, considering the 10 genes with the highest z-score for membership of these gene sets, we find that the genes implied by the joint analysis have a higher likelihood of gene set membership (see the DEPICT paper for a detailed explanation [33]), thus resulting in more coherent gene sets. For instance, the joint analysis identifies the LDLR (LDL receptor) gene, which has a high membership likelihood for the REACTOME “metabolism of lipids and lipoproteins” gene set. The pairwise analysis does not identify LDLR, making this gene set less enriched. These results show that joint pleiotropy analysis of multiple traits yields more biologically relevant insights compared to pairwise analysis of those same traits.

### Runtime increases marginally with the number of traits analyzed

To assess how the runtime scales with the number of traits analyzed, we picked all traits that were affected by the most pleiotropic SNP, rs495828: AS, BC, CAD, CES, DBP, HDL, HF, IS, LDL, T2D, TAGC, and TC. In this order, we picked the first p traits and timed PolarMorphism (see Figure 5). Runtime increases slightly with larger p, but the effect is small. There is a large difference between p = 2 and p > 2 because we use different approaches if p > 2 (see methods).

### Comparison with other methods

To compare PolarMorphism to existing methods, we ran: PolarMorphism, intersection, PLACO, and PRIMO on a selection of traits (IS and myocardial infarction). We compared the individual SNPs and loci that were identified as pleiotropic by each method. Four loci are found by all methods. Intersection does not identify more than those four loci. PLACO and PolarMorphism both find 21 loci (19 of which are identical), PRIMO finds 13 loci that were also identified by PLACO and PolarMorphism. PLACO and PolarMorphism use a fundamentally different approach to identify pleiotropy: whereas PLACO tests if the effect for both traits is not equal to zero, PolarMorphism first tests whether the overall effect (distance) is different than expected and then tests the sharedness of a SNP.

We timed each method from cleaned input data (already in memory, timing done in R) to results. The number of pleiotropic loci that were found by each method and the speed of generating results (in number of input SNPs per second) are provided in table 2. These data show that PLACO does not identify more loci than PolarMorphism and is slower.

**Table 2.** Comparison of methods. HOPS and PLEIO were not run because they use a pleiotropy definition that includes single-trait SNPs.

|  | Max p | Decorrelation? | # of pleiotropic loci found | speed (1k SNPs/second) |
| --- | --- | --- | --- | --- |
| PolarMorphism | - | Yes | 21 | 63 |
| PLACO | 2 | Yes | 21 | 0.61 |
| Primo | - | No | 13 | 86 |
| HOPS | - | Yes | - | - |
| PLEIO | - | No | - | - |

## Discussion

We have developed a new method that identifies pleiotropic SNPs with an effect on multiple traits. PolarMorphism can be used on combinations of two or more traits. It uses GWAS summary statistics and corrects for correlation in effect sizes arising from genetic correlation or potential sample overlap. The potential applications of PolarMorphism include a) identifying SNPs that are shared between traits within a trait domain to learn more about the domain-wide biological processes, b) identifying SNPs that are shared among a diverse set of traits to find general biological processes and c) using the identified SNPs to inform new trait ontologies. As an example, we apply PolarMorphism to a set of traits from different domains.

The network analyses indicate that there are no trait domains that only share SNPs within the domain. We observe that most SNPs are associated with traits within one or across two trait domains. We zoomed in on the stroke domain, which has very little domain-specific SNPs. This may mean that the stroke traits are associated with general SNPs or that the stroke traits do not share many biological pathways. Each ischemic stroke subtype shares more SNPs with non-stroke traits than with the other ischemic stroke subtypes. Note that these networks are heavily influenced by the choice of included traits. Conclusions drawn about the networks in this study are therefore not necessarily general, as each trait could share SNPs with a number of traits that were not included.

We compared PolarMorphism with similar methods. PolarMorphism identifies more pleiotropic SNPs than the standard intersection method and than PRIMO. PLACO identifies the same number of pleiotropic loci as PolarMorphism. However, PolarMorphism finished analysis of 1 million SNPs in less than 20 seconds (compared to >25 minutes for PLACO), making analysis of many trait combinations feasible. Furthermore, PLACO can only be used to analyze two traits together while PolarMorphism can analyze a theoretically unlimited number of traits. A five-fold increase in the number of identified pleiotropic loci for the lipid domain indicates that analyzing more than two traits is much more powerful than combined results from the respective pairwise analyses.

## Methods

### PolarMorphism for two traits

PolarMorphism works on uncorrelated, standardized data. *z_x_* and *z_y_* are vectors of length *m* containing the z-scores of SNPs 1 to *m* for trait x and trait y, respectively. We calculate polar coordinates r and θ: r is the distance from the origin, and θ is the angle of the vector from the origin to the point (z_x_, z_y_). 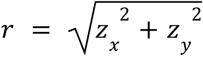 and 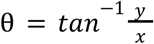.

We first test whether *r* comes from a central chi distribution with degrees of freedom equal to the number of traits *p*. The chi distribution describes the distribution of the square root of the sum of squared normally distributed variables. The distribution of p-values from this test is used to calculate q-values, which are FDR-corrected p-values [34]. For all SNPs that have an effect, we want to know whether that effect is shared. We perform a four-fold transform of θ that ‘folds’ all quadrants of the Cartesian plane on top of each other and stretches it to make sure the angles can take any value on the circle [35]: *θ_trans_* = 4θ *modulo* 2π. The von Mises distribution describes angular data. It takes into account that θ = 0 is equal to θ = 2*π*. It has two parameters: *θ_mu_* is the mean value, and kappa (*κ*) is a concentration parameter that is similar to the inverse of the variance. *θ_mu_* is zero under the null hypothesis of trait-specific effect. See the Supplementary methods for a description of how we obtained estimates for *κ*. Using the distribution of the observed *r* p-values for the distances of all SNPs, and the fact that p-values follow a uniform distribution under the null hypothesis, the false discovery rate (FDR) for each SNP can be calculated. This q-value gives the FDR if this SNP and all SNPs with a lower p-value would be called significant. We keep the SNPs that show a significant overall effect (r q-value < 0.05) and use the distribution of observed θ p-values for these SNPs to calculate θ q-values. We filter on *θ* q-value < 0.05 to obtain SNPs that are significantly shared (FDR < 0.05).

### PolarMorphism for more than two traits

#### Converting Cartesian to hyperspherical coordinates

The distance of a SNP *i* in more than two dimensions is a straightforward extension of the distance in two dimensions:

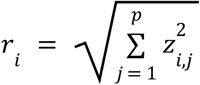

Where *z_i,j_* is the z-score of SNP *i* for trait *j*. Describing the orientation of a SNP for *p* traits involves calculating the corresponding *p*-dimensional hyperspherical coordinates. This gives an additional angle for each added trait. Fortunately, this problem can be simplified. We define 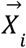 as the vector from the origin of the p-dimensional sphere to an observed SNP, and 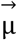 as the vector from the origin to the expected position of the SNP under the null hypothesis of trait-specific effect, along one of the axes. The goal is to determine the angular difference between 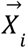 and 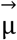. We choose 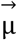 such that it lies along the axis that is closest to 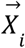. In other words, we construct 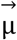 as a vector with zeros for each coordinate except for the coordinate with the highest absolute value for the SNP under consideration. We set the length of 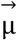 equal to the length of 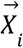 (the distance *r*), so the only non-zero value of 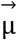 is set to *r*. The two vectors of interest always lie in a 2-dimensional plane, regardless of the number of traits p. The dot product of the vectors is a scalar and is equal to:

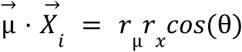

therefore

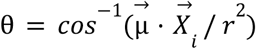

Which can be rewritten as

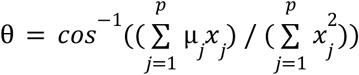

This angle should be normalized so the maximum value is always *π*, regardless of *p*. The angle is maximal if all coordinates of a SNP have the same value (which we will call *x*). Recall that 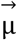 has zeros for all coordinates but one. If θ is maximal, we can rewrite the expression for θ as:

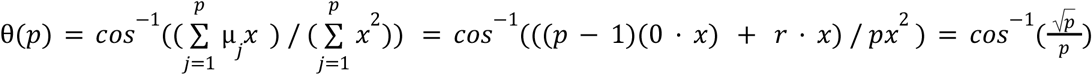

The final correction factor with which the angles should be multiplied can then be obtained by dividing *2π* by the result of this formula.

#### Testing the significance of *r and θ*

To test the significance of r, we use the same procedure as for two traits. In this case the degrees of freedom is equal to the number of traits *p*. To assign significance levels to the angle *θ*, we use the von Mises-Fisher distribution, which is an extension of the von Mises distribution. The probability density function of the von Mises Fisher distribution is given by:

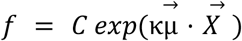

Where C is a normalization constant, *κ* is the concentration parameter, 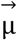 is the unit vector of the expected direction and 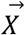 is the observed unit vector (i.e. the vector of the SNP divided by its length to get unit length). The inner product 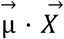 can be rewritten as cos(θ), where θ is the angle between the expected and observed vectors:

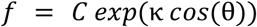

Functions to obtain the probability density function and the normalization constant *C* are implemented in the vMF package in R [36]. To obtain a cumulative density function the probability density function needs to be integrated. The definite integral for *exp*(*κ cos*(θ)) can not be defined using elementary functions. However, the exponent has the following series representation:

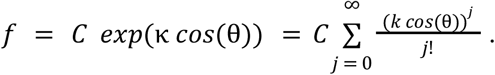

The integral is then equal to:

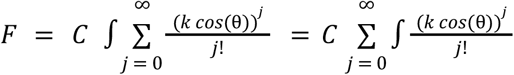

The term (as a function of the iterator j) does have an indefinite integral:

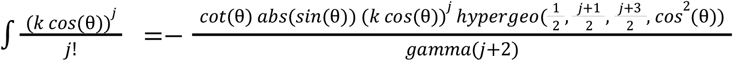

where *cot* is the cotangent function, *hypergeo* is the hypergeometric function and *gamma* is the gamma function. We implemented the summation so that it stops when the last added term is smaller than a user-defined value (called ‘tol’ in our R package). We use the hypergeo package for the hypergeometric function [37]. The values for *κ* as a function of *p* that we derived for p = 2 still apply here, because θ still describes a two-dimensional angle.

### Preprocessing the summary statistics

We used publicly available summary statistics for the 41 traits shown in table 1. Data were obtained from the sources provided in Supplemental table 2, which also contains references to the respective papers they were described in. We aligned reference and alternative allele across all traits, and filtered using the list of high-confidence SNPs provided with the LDSC software.[23] We divide effect sizes by their standard error to obtain z-scores. We calculate the covariance matrix on the subset of SNPs that do not have a large overall effect. To this end, the covariance is calculated only on SNPs that have a mahalanobis distance smaller than 5. We use the ZCA-cor whitening method in the ‘whitening’ package in R [38], to decorrelate the data while ensuring that the x and y components of the transformed z-scores maximally correlate with the x and y components of the original z-scores.

### Inferring relationships between traits from pleiotropic SNPs

For all trait pairs, we ran PolarMorphism and clumped the significant SNPs with Plink, using the q-values instead of p-values (--clump-kb 5000000, --clump-p1 0.05, --clump-p2 0.05, --clump-r2 0.2) [39]. We make an adjacency matrix from the number of shared loci per trait combination and use this to construct a graph using the igraph package in R [40]. We did the same per SNP to obtain SNP-specific networks. To create domain networks from the trait networks we draw an edge between domain A and B if an edge exists between any trait of domain A and any trait of domain B.

### Gene set enrichment analysis in DEPICT

We changed the following settings from the default: association_pvalue_cutoff: 0.05 to accommodate for the fact that we use q-values instead of p-values. We performed gene set enrichment using the default gene sets provided by the DEPICT authors, but only considered gene sets from gene ontology [41], REACTOME [42], KEGG[43] and the PPI networks as defined by the DEPICT authors using the InWeb database [44] for further analysis.

### Inferring relationships between traits from genetic correlation

To infer relationships between traits from genetic correlation, we ran LDSC[23] using the GenomicSEM [45] package in R. We calculated p-values from the correlation coefficients and their standard errors using the pnorm function in R, and used a Bonferroni corrected p-value threshold of 6.4*10^-5^ to correct for 780 trait combinations tested. For this purpose, we made an adjacency matrix from the genetic correlation for each trait combination and used this to make a graph using the igraph package in R.[40]

### Comparison with other methods

Intersection refers to the straight-forward approach of finding shared SNPs: take the intersection of the SNPs that were significant for trait X and those that were significant for trait Y. We used the R package for HOPS (HOrizontal Pleiotropy Score) [14] We used our pre-processed z-scores (whitened). We ran HOPS both with and without polygenicity correction and used only the Pm p-values. We used the command line tool written in Python for PLEIO (Pleiotropic Locus Exploration and Interpretation using Optimal test) [15]. We used z-scores (not whitened and not corrected for LD-score) and supplied the sample sizes of the original GWAS. We used the R package for PRIMO (Package in R for Integrative Multi-Omics association analysis) [16]. We used PRIMO based on p-values. For the alt_props parameter (the expected proportion of SNPs that follow the alternative hypothesis per trait) we supplied the proportion of SNPs that were significant for trait 1 (q-value < 0.05) over all SNPs, idem for trait 2 (q-value < 0.05). We supplied c(2,2) for the dfs parameter. We used the R package for PLEIO (pleiotropic analysis under composite null hypothesis) [15]. We used whitened z-scores (not corrected for LD-score). We used the VarZ function to calculate the covariance matrix and supplied that, with the z-scores, to the placo function.

To assess how many loci were found by each method, we LD-pruned the significantly shared SNPs. For each method and for each locus, we checked if any of the SNPs in that locus were also found by another method. If that was the case, we gave that locus the same identifier in each method. Afterwards, we determined the loci that were found by all methods and those that were found by only one or a subset of the methods. We ran Intersection, HOPS (with polyenicity correction), PRIMO, PLACO, and PolarMorphism on the same data while supplying a dataframe with an increasing number of rows. For the Intersection method we added q-value calculation from the original GWAS p-values and a filtering step on both q-values to make it a fair comparison with the other methods. All five methods are written in R, therefore we timed them in R using the tictoc package [46]. Running the software in the terminal could have a different runtime, but this does allow us to compare the runtimes among the methods.

## Data availability

The PolarMorphism results presented in this paper are available on Zenodo, at https://dx.doi.org/10.5281/zenodo.5844193.

## Acknowledgements

The authors thank René Eijkemans for helpful discussions. JvB is supported by R01NS100178 from the National Institute of Health. JdR is supported by a Vidi Fellowship (639.072.715) from the Dutch Organization for Scientific Research (Nederlandse Organisatie voor Wetenschappelijk Onderzoek, NWO). SWvdL is funded through grants from the Netherlands CardioVascular Research Initiative of the Netherlands Heart Foundation (CVON 2011/B019 and CVON 2017-20: Generating the best evidence-based pharmaceutical targets for atherosclerosis [GENIUS I&II]). We are thankful for the support of the ERA-CVD program ‘druggable-MI-targets’ (grant number: 01KL1802), the EU H2020 TO_AITION (grant number: 848146), and the Leducq Fondation ‘PlaqOmics’.

## Notes

https://github.com/UMCUGenetics/PolarMorphism

https://dx.doi.org/10.5281/zenodo.5844193

### Competing Interest Statement

The authors have declared no competing interest.

