## Supplemental_Information for "PolarMorphism enables discovery of shared genetic variants across multiple traits from GWAS summary statistics": PolarMorphism_supplemental.docx

### Table S1. Variables and their definitions as used in this paper

|  |  | **description** | **definition** | **distribution** |
| --- | --- | --- | --- | --- |
| **Variables** | N | sample size | number of observations |  |
|  | m | number of SNPs |  |  |
|  | p | number of traits |  |  |
|  | X | effect size on trait x |  | $X \sim N(\mu_{x}, {\sigma_{x}}^{2})$ |
|  | Y | effect size on trait y |  | $Y \sim N(\mu_{y}, {\sigma_{y}}^{2})$ |
|  | μ | mean |  |  |
|  | σ | standard deviation |  |  |
|  | r | distance from the origin | $r = \sqrt{x^{2}+y^{2}}$ | $r \sim\chi(p, \lambda)$ |
|  | θ | angle with the x-axis | $\theta=tan^{-1}(\frac{y}{x})$ | $\theta\sim vonMises(\mu_{\theta}, \kappa)$ |
|  | $\theta_{trans}$ | fourfold transform of θ | $\theta_{trans}:=4\theta modulo 2\pi$ | $\theta_{trans}\sim vonMises(0, \kappa)$ |
|  | $\kappa$ | Concentration parameter |  |  |
|  | *z_max_* | Maximum of absolute values of X and Y for a SNP |  |  |
| **Subscripts** | x | of trait x |  |  |
|  | y | of trait y |  |  |
|  | θ | of θ |  |  |
|  | obs | observed |  |  |
| **Operators** | E[X] | expected value/expectation of X |  |  |
|  | var[X] | variance of X |  |  |
|  | e^x^ | the (natural) exponential function | $e^{x}:=\sum_{k = 0}^{\infty} \frac{x^{k}}{k!}$ |  |
|  | exp(x) | Alternate notation of e^x^ |  |  |

#### Table S2. GWAS used

| **Trait or disease** | **Abbreviation** | **Sample size** | **Consortium** | **PMID** | **Reference** | **Ancestry** |
| --- | --- | --- | --- | --- | --- | --- |
| Alzheimer's Disease | AD | 455258 |  | 30617256 | Jansen et al., 2019 [[47]](https://paperpile.com/c/kggNEW/kNUQ) | European |
| Atrial Fibrillation | AF | 588190 |  | 29892015 | Roselli et al., 2018 [[48]](https://paperpile.com/c/kggNEW/xh5j) | Multi ancestry |
| Amyotrophic Lateral Sclerosis | ALS | 43259 |  | 27455348 | van Rheenen et al., 2016 [[49]](https://paperpile.com/c/kggNEW/R8YI) | European |
| Any stroke | AS | 446696 | MEGASTROKE | 29531354 | Malik et al., 2018 [[28]](https://paperpile.com/c/kggNEW/LurS) | European |
| Autism Spectrum Disorder | ASD | 46350 | PGC | 30804558 | Grove et al., 2019 [[50]](https://paperpile.com/c/kggNEW/UfkA) | European |
| Asthma | Asthma | 142486 |  | 29273806 | Demenais et al., 2018 [[51]](https://paperpile.com/c/kggNEW/iTEk) | Multi ancestry |
| Breast Cancer | BC | 256123 |  | 29059683 | Michailidou et al., 2017 [[52]](https://paperpile.com/c/kggNEW/Rj2c) | Multi ancestry |
| Bipolar Disorder | BIP | 198882 | PGC | 31043756 | Stahl et al., 2019[[53]](https://paperpile.com/c/kggNEW/wE51) | European |
| Body Mass Index | BMI | 693529 | GIANT | 30124842 | Yengo et al., 2018 [[54]](https://paperpile.com/c/kggNEW/BHbn) | European |
| Coronary Calcification | CAC | 15523 |  | 23561647 | Van Setten et al. 2013 [[55]](https://paperpile.com/c/kggNEW/zbfm) | European |
| Coronary Artery Disease | CAD | 154654 | CARDIoGRAMplusC4D+UKBB | 28714975 | Nelson et. al, 2017 [[56]](https://paperpile.com/c/kggNEW/5kC5) | European |
| Cardio-Embolic stroke | CES | 521612 | MEGASTROKE | 29531354 | Malik et al., 2019 [[28]](https://paperpile.com/c/kggNEW/LurS) | European |
| Carotid Intima-Media Thickness | cIMT | 71128 | CHARGE | 30510157 | Franceschini et al. 2018 [[57]](https://paperpile.com/c/kggNEW/hUob) | Multi ancestry |
| Cigarettes per Day | CpD | 74035 | TAG | 20418890 | The Tobacco and Genetics Consortium, 2010 [[58]](https://paperpile.com/c/kggNEW/sn9j) | European |
| Diastolic Blood Pressure | DBP | 1050906 |  | 30224653 | Evangelou et al., 2018 [[59]](https://paperpile.com/c/kggNEW/btwR) | European |
| Depressive symptoms | DS | 298420 |  | 27089181 | Okbay et al., 2016 [[60]](https://paperpile.com/c/kggNEW/Rs4S) | European |
| Educational Attainment | EA | 293723 |  | 27225129 | Okbay et al., 2016 [[61]](https://paperpile.com/c/kggNEW/IeyF) | European |
| Ever smoked | EvrSmk | 74035 | TAG | 20418890 | The Tobacco and Genetics Consortium, 2010 [[58]](https://paperpile.com/c/kggNEW/sn9j) | European |
| Forearm Bone Mass Density | FABMD | 53236 | GEFOS | 26367794 | Zheng et al., 2015 [[62]](https://paperpile.com/c/kggNEW/LMoa) | European |
| Femoral Neck Bone Mass Density | FNBMD | 53236 | GEFOS | 26367794 | Zheng et al., 2015 [[62]](https://paperpile.com/c/kggNEW/LMoa) | European |
| Former Smoker | FrmrSmk | 74035 | TAG | 20418890 | The Tobacco and Genetics Consortium, 2010 [[58]](https://paperpile.com/c/kggNEW/sn9j) | European |
| High-Density Lipoprotein | HDL | 1888577 | GLGC | 24097068 | Global lipids Genetics Consortium , 2013 [[63]](https://paperpile.com/c/kggNEW/cHIS) | European |
| Height | Height | 693529 | GIANT | 30124842 | Yengo et al., 2018 [[54]](https://paperpile.com/c/kggNEW/BHbn) | European |
| Heart Failure | HF | 488010 |  | 30586722 | Aragam et al. 2018 [[64]](https://paperpile.com/c/kggNEW/YfAU) | European |
| Inflammatory Bowel Disease | IBD | 86640 |  | 26192919 | Liu et al., 2015 [[65]](https://paperpile.com/c/kggNEW/ttrh) | Multi ancestry |
| Insomnia | Insomnia | 386533 |  | 30804565 | Jansen et al., 2018 [[66]](https://paperpile.com/c/kggNEW/Dbtp) | European |
| Intelligence Quotient | IQ | 269867 |  | 29942086 | Savage et al., 2018 [[67]](https://paperpile.com/c/kggNEW/DkBg) | European |
| Any Ischemic stroke | IS | 446696 | MEGASTROKE | 29531354 | Malik et al., 2019 [[28]](https://paperpile.com/c/kggNEW/LurS) | European |
| Large Artery Stroke | LAS | 446696 | MEGASTROKE | 29531354 | Malik et al., 2019 [[28]](https://paperpile.com/c/kggNEW/LurS) | European |
| Low-Density Lipoprotein | LDL | 1888577 | GLGC | 24097068 | Global lipids Genetics Consortium , 2013 [[63]](https://paperpile.com/c/kggNEW/cHIS) | European |
| Onset Smoking | logOnset | 74035 | TAG | 20418890 | The Tobacco and Genetics Consortium, 2010 [[58]](https://paperpile.com/c/kggNEW/sn9j) | European |
| Lumbar Spine Bone Mass Density | LSBMD | 53236 | GEFOS | 26367794 | Zheng et al., 2015 [[62]](https://paperpile.com/c/kggNEW/LMoa) | European |
| Major Depression Disorder | MDD | 480359 | PGC | 29700475 | Wray et al., 2018 [[68]](https://paperpile.com/c/kggNEW/duZI) | European |
| Neuroticism | Neuroticism | 298420 |  | 27089181 | Okbay et al., 2016 [[60]](https://paperpile.com/c/kggNEW/Rs4S) | European |
| Nonischemic Cardiomyopathy | NICM | 488010 |  | 30586722 | Aragam et al. 2018 [[64]](https://paperpile.com/c/kggNEW/YfAU) | European |
| Parkinson's Disease | PD | 1456306 |  | 31701892 | Nalls et al.,2019 [[69]](https://paperpile.com/c/kggNEW/Gift) | European |
| Plaque Presence | Plaque | 48434 | CHARGE | 30510157 | Franceschini et al. 2018 [[57]](https://paperpile.com/c/kggNEW/hUob) | Multi ancestry |
| Pulse Pressure | PP | 1050906 |  | 30224653 | Evangelou et al., 2018 [[59]](https://paperpile.com/c/kggNEW/btwR) | European |
| Prostate Cancer | PrCa | 140306 |  | 29892016 | Schumacher et al., 2018 [[70]](https://paperpile.com/c/kggNEW/rY6I) | European |
| Systolic Blood Pressure | SBP | 1050906 |  | 30224653 | Evangelou et al,. 2018 [[59]](https://paperpile.com/c/kggNEW/btwR) | European |
| Small Vessel Disease | SVD | 446696 | MEGASTROKE | 29531354 | Malik et al., 2019 [[28]](https://paperpile.com/c/kggNEW/LurS) | European |
| Subjective Well-Being | SWB | 298420 |  | 27089181 | Okbay et al., 2016 [[60]](https://paperpile.com/c/kggNEW/Rs4S) | European |
| Type 2 Diabetes | T2D | 898130 | DIAGRAM | 30297969 | Mahajan et al., 2018 [[71]](https://paperpile.com/c/kggNEW/j56n) | European |
| Type 2 Diabetes adjusted for BMI | T2DadjBMI | 898130 | DIAGRAM | 30297969 | Mahajan et al., 2018 [[71]](https://paperpile.com/c/kggNEW/j56n) | European |
| Total Cholesterol | TC | 1888577 | GLGC | 24097068 | Global lipids Genetics Consortium , 2013 [[63]](https://paperpile.com/c/kggNEW/cHIS) | European |
| Triglycerides | TG | 1888577 | GLGC | 24097068 | Global lipids Genetics Consortium , 2013 [[63]](https://paperpile.com/c/kggNEW/cHIS) | European |

#### Table S4. Significant loci of pairwise analyses of HDL, LDL, TC and TG

#### Table S5. Significant gene sets of pairwise analyses of HDL, LDL, TC and TG

#### Table S6. Significant loci of joined analysis of HDL, LDL, TC and TG

#### Table S7. Significant gene sets for joined analysis of HDL, LDL, TC and TG

#### Table S8. The number of single domain SNPs, which are only associated with traits within the domain in question.

#### Supplementary text 1

Estimating the concentration parameter of the von Mises distribution

Intuitively, the expected distribution of the angle θ is more concentrated if a SNP is further away from the origin, because the same variation in z_x_ and z_y_ translates to a smaller variation in θ. This indicates that there should be a relationship between *r* and *𝜅*.

Given a sample of angles θ coming from a von Mises distribution with a certain *θ_mu_* and 𝜅, the values of both parameters can be estimated from the *resultant vector* of the data [[72]](https://paperpile.com/c/kggNEW/kCel). The resultant vector *R* is the vector sum of all observed angles θ in the sample, if each angle is a vector with unit length and angle θ with the x-axis. The angle of *R* with the x-axis is then equal to *θ_mu_*. The length of the *mean resultant vector* $\underline{R}$ - *R* divided by the number of angles - says something about the spread in the data. If the variation in θ is large, $\underline{R}$ is small and if the angles are very concentrated around *θ_mu_*, $\underline{R}$ is large. The length of $\underline{R}$ can take on values from 0 (if the angles are maximally spread out and each individual observation is canceled out by an angle in the opposite direction) to 1 (if the angles are all identical to *θ_mu_*). If $\left| \underline{R} \right|$ (the length of $\underline{R}$) is 0, 𝜅 is 0, and if $\left| \underline{R} \right|$ is 1, 𝜅 is equal to infinity. An analytical expression for the relationship between *r* and $\left| \underline{R} \right|$ could not be found because it depends on the expected value of the inverse distribution, which does not exist. An empirical relationship was found by simulating bivariate normal distributions with bivariate mean (*r*, 0) for increasing values of *r*, from 0 until 50. We used the mle.vonmises function from the circular package [[73]](https://paperpile.com/c/kggNEW/rA8v) to calculate 𝜅 for each simulated value of r. We include a table with values of r and corresponding simulated values of 𝜅 with our R package. See figure 6 for the empirical relationship between r and 𝜅.

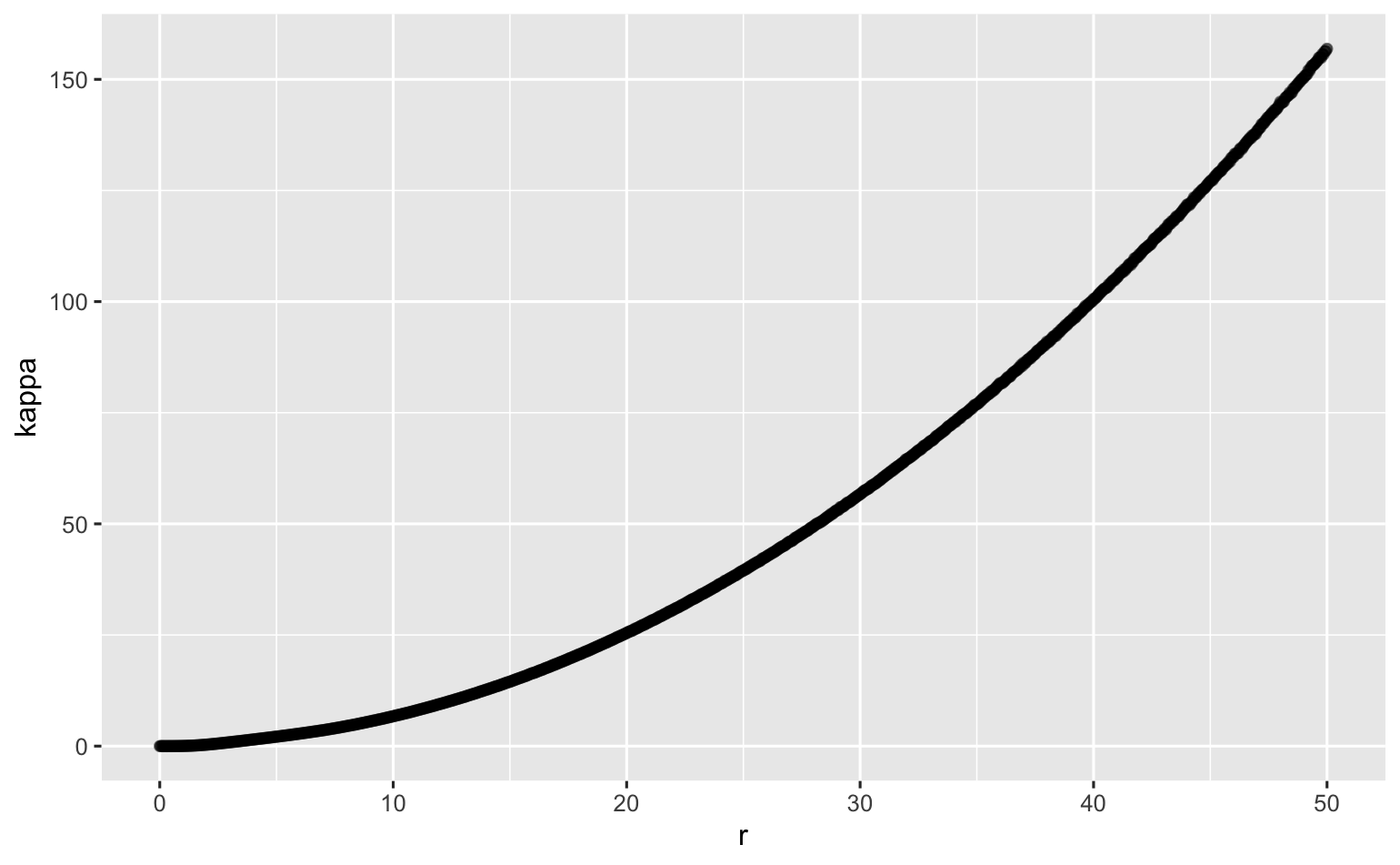

*Figure 6. Visualization of the relationship between kappa and r*
